# Visuomotor phase-locked loop reproduces elliptic hand trajectories across different rhythms

**DOI:** 10.1101/2022.07.20.500761

**Authors:** Adam Matić

**Author notes:** ^*^Correspondence.

## Abstract

A well-known phenomenon in human hand movement is the correlation between speed and curvature, also known as the speed-curvature power law (V≈kC^β^). In drawing elliptic shapes, the exponent is often found to be β ≈ −1/3, however it is not clear why the power law appears and why the exponent is near −1/3. More fundamentally, it is not clear how do people track elliptic targets. In answering these questions, I’ve analyzed trajectories of participants’ cursors while they tracked visual targets moving along elliptical paths, across different target speed profiles and cycle frequencies. The speed-curvature power law emerged when drawing ellipses at about 1 Hz or faster, regardless of the target speed profile, and it did not emerge for lower frequency movements. Analysis of the position frequency spectrum shows that the target-cursor trajectory transformation may be seen as a low-pass filter. Comparison of different hypothetical salient features of the visual field shows that phase difference (angular difference between the cursor and the target) and size difference (difference in the sizes of the elliptic paths) are the features most likely used in the task. The next experiment confirmed that phase and size difference could be controlled variables because participants kept them stable even under direct pseudorandom disturbances. A numerical model simulating the sensorimotor processes of the participant, similar to a phase-locked loop, using the visual features of phase and size difference as controlled variables, performed the same target tracking tasks as the participants. When fitted, the model closely replicated position and speed profiles of the participants across all trials, as well as the emergence of the power law at high frequencies. The model also reproduced the trajectories of participants in the experiment with direct pseudorandom disturbances. In conclusion (1) the speed-curvature power law emerges as a side effect of movement system properties, namely low-pass filtering in the sensorimotor loop; (2) people could be tracking elliptical targets by varying the frequency and amplitude of an internal pattern generator until the produced phase and shape size match the target’s phase and shape size. The model generates new hypotheses about the neural mechanisms of rhythmic movement control.

## Introduction

Hand movement is created in the continuous and closed loop interaction between the brain, body and environment. An often-studied phenomenon in hand movement is the correlation between speed and curvature, first noticed by Binet and Courtier (1893) and Jack (1895) in handwriting as slower movements of the pen in areas of high curvature, and faster movements of the pen in areas of low curvature. The phenomenon was later studied by Viviani and Terzuolo (1982) and formalized as the ‘speed-curvature power law’ by Lacquaniti et al (1983). The name “power law” comes from the empirical relationship V≈kC^β^ where speed V is approximately equal to curvature C raised to the power β, times a constant k. Speed, or tangential speed, is defined as the magnitude of the velocity vector, 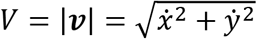 and curvature as the reciprocal of the radius if the osculating circle, 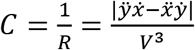, where x and y are coordinates of the point P in the plane (Figure 1A). In rhythmical elliptic movement, the exponent β is often found to be −1/3 (Figure 1B). The value of the exponent β expresses the degree of “slowing down in curves”: for β=0, there is no slowing, the speed is constant (Figure 1C); for β=−1/3, the speed is slightly lower in curves than in straight parts, and for β=−2/3, the speed is much lower in curves than in straight parts.

**Figure 1.**
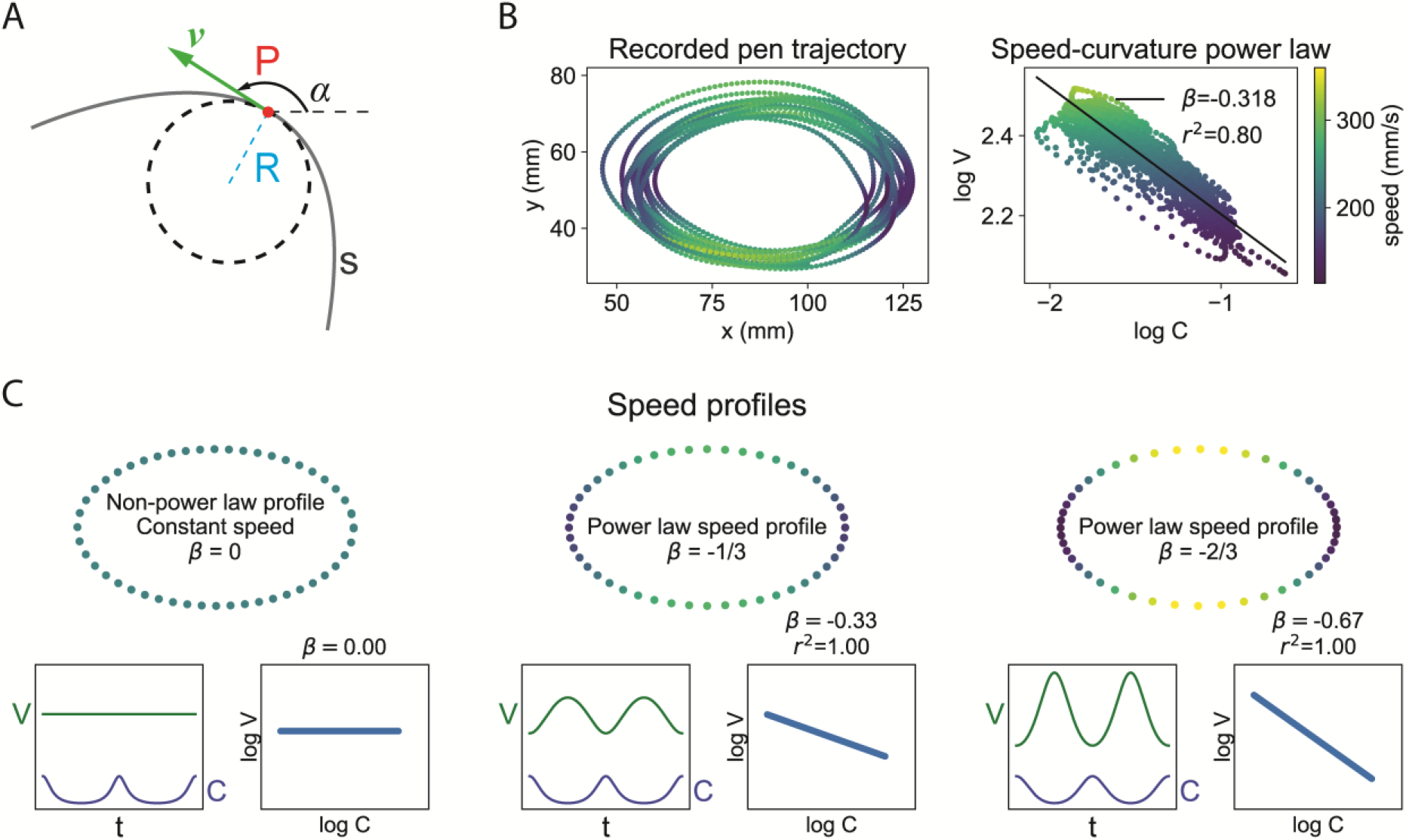
The hand movement speed-curvature power law, V≈kC^β^, or log V ≈ log k + β*log C. **A**) Geometric and kinematic definitions of the variables: the point P moves along the curve s; at each instant of time, its velocity can be described by the vector **v**, speed V is the magnitude of velocity, and curvature C is the reciprocal of the radius R of the osculating circle (C = 1/R). The direction of movement with respect to the x-axis is **α**, and its first time-derivative is the angular velocity A. The variable k is a constant related to average speed. **B)** An example of an empirical trajectory conforming to a speed-curvature power law: a participant moves the pen faster (bright green) in areas of low curvature and slower (dark green) in areas of high curvature. On a log-log plot, the relationship between speed and curvature is linear, with the slope β, intercept log k and coefficient of determination r^2^ **C**) Examples of the speed-curvature relationships for equal curvature profiles, but different speed profiles. In the first case, the speed is constant, and there is no power law. In the second case, the speed is slightly lower in areas of high curvature, with the exponent β= −1/3 (~0.33) and r^2^=1. This profile is often found in elliptic human hand movements. In the third case, the speed is much lower in areas of high curvature than in areas of low curvature, with the exponent β= −2/3 (~0.67) and r^2^=1.

An alternative form of the law uses angular speed, defined as the rate of change of the direction of the velocity vector, instead of tangential speed. The two forms have equivalent exponents, however, angular speed tends to have much higher correlations to curvature than tangential speed, as well as higher coefficients of determination in the regression estimate of the power law, and should be avoided when estimating the speed-curvature power law (Matić and Gomez-Marin, 2022). In further analysis, I will be using tangential speed and curvature.

Numerous studies found support for the law, from manually tracking visual targets (Viviani and Mounoud, 1990), drawing ellipses in water (Catavittelo et al, 2017), walking trajectories (Vieilledent et al, 2001) to changes in the exponent during child development (Viviani and Schneider, 1991). When drawing different shapes, the exponent has a spectrum of values different from −1/3 (Huh and Sejnowski, 2015). The power law was also found in movements of hands of monkeys and in population codes in their motor cortex (Schwartz, 1994), as well as in movements of the drosophila larvae (Zago et al. 2016). For a recent review of the evidence for the speed-curvature power law, the statistics used and hypotheses of its origin, see Zago, Matić et al. (2017).

Hypotheses aiming to explanation the emergence of the power law may be divided into two broad groups - central origin theories and emergence theories. The central origin group argues that, since the speed-curvature relationship is found in so many different types of movements, it is likely a part of the planning strategy for every movement. For example, it could be a result of minimizing jerk (Viviani and Flash 1995), maximizing smoothness (Huh and Sejnowski, 2015) or minimizing endpoint variance (Harris and Wolpert, 1998). The second group of explanations favors emergence of the power law in the interaction of the brain, body with the environment, mainly due to low pass filtering (Gribble and Ostry 1996; Schaal and Sternad, 2001).

The relationship of average speed or rhythm of drawing and the exponent of the power law has not been explored in detail, although Lacquaniti et al (1983, Figure 4) reported different values of the exponent for three different rhythms of drawing, and Wann et al (1988) reported two different exponents for two different rhythms, all of relatively fast movement.

Here, I explore target tracking over a wider range of rhythms for a constant size elliptic trajectory, and the relationship between the rhythm and the power law exponent and the coefficient of determination. Next, I estimate salient visual features used by the participant in the task, and build a numerical model, based on the findings, that performs the same task.

My basic assumption was that during the task, participants observe and maintain (control) certain visual variables at their respective reference values. It was not obvious what these variables were, and how to describe them mathematically. The deceptively simple strategy, suggested by Powers (1973, 1978) is to find variables that remain stable despite being disturbed by the experimenter.

The search starts by generating precise mathematical definitions of the hypothetical controlled variables. This definition needs to make clear what are the effects of the experimenter-generated disturbances on the controlled variable, and the effects of participant-generated behavior on the controlled variable. For instance, the controlled variable may be defined as a distance between the target and the cursor, the disturbance is, therefore, the position of the target, and the behavior of the participant is the position of the cursor. Both the target and the cursor can affect the controlled variable in a precisely defined way. The experimenter can directly perturb this hypothetical controlled variable by generating a target trajectory and then observe corrective actions by the participant; or alternatively the experimenter can insert a disturbance between the pen and the cursor on the screen.

If the hypothetical controlled variable is stable relative to the variance of the perturbation, it might be a good approximation for the variable controlled by the participant. If it is not stable, there are several possibilities – (1) the variable is not controlled, (2) the variable is controlled, but the bandwidth of the disturbance is too wide (the task is too difficult), (3), the variable is controlled, but the reference level was not stable, etc. The task should be designed to be not too difficult, and the reference level should be stable, or with a known variation.

Aside from the stability of the controlled variable, another useful statistic is a *low* correlation between the potential controlled variable and the disturbance variable – meaning that a controlled variable is unaffected by the disturbance, and as a consequence it is uncorrelated to it.

When a sufficiently good and precise definition of the hypothetical controlled variable is found, then a generative numerical model, a simulation of the participants’ sensorimotor loop in the can be built, and fitted to each participant individually. For further validation, the behavior of the model can be compared to the behavior of the participants in new tasks where the model was not fitted. Good performance of the model in new tasks would support the explanatory and predictive power of the model and the conclusion that the mathematical definition of the controlled variable is a good approximation of the variable controlled by the participant. There is always a possibility of finding a better approximation.

The model should be made as computationally simple as possible in order to be biologically plausible. Signals generated by the numerical model may be direct correlates of neural signals in the nervous system of the participant, and this might be verified in an independent experiment.

Strong invariances in participants’ movements, such as the speed-curvature power law, may be used to verify a model of rhythmic behavior: the power law needs to appear in the same conditions for the model as for the participants, and also it needs to *not* appear in the same conditions for the model, where it does not appear for the participants.

### Data and code availability

The data collected in this research, along with jupyter notebooks and python code used to analyze the data and prepare the figures is available at https://github.com/adam-matic/visuomotor-phase-locked-loop

#### Experiment 1: Tracking targets along rhythmic elliptical trajectories

##### Method

In the first experiment, participants tracked a target with a cursor on the computer monitor. Participants (N=3, male, right-handed, age 30-36, including the author of the paper) were seated, looking at the computer monitor showing a cursor and a target. They were holding an electronic pen in their dominant hand, pen positioned on the graphics tablet (Wacom Intous PTH S). They were instructed to keep the cursor and target as close as possible. Target trajectories had three different speed profiles: (1) β=0, constant speed, (2) β=−1/3, ‘natural’ speed with slight slowing in the curved areas, and (3) β=−2/3, excessive slowing in the curved areas, (see Figure 1C). All three profiles were generated for nine fundamental frequencies of the target: 0.27, 0.40, 0.54, 0.67, 0.81, 0.94, 1.07, 1.21, and 1.34Hz corresponding to average speeds of the target: 44, 67, 91, 113, 135, 158, 180, 203, 227 mm/s. The target always moved in a counter-clockwise direction. The trials were presented in a random order. Each trial started when the participant pressed the space key, and lasted for 20 seconds, after which the participant could rest. The monitor displayed the target and cursor at 60Hz, while the tablet recorded the pen position data at 200Hz.

Target trajectories were formed by first generating a high-resolution elliptical path, and then rescaling the temporal distances between the points so that the speed conformed to a desired speed-curvature power law. The resulting target trajectory was splined and resampled to have the points equally spaced in time, and rescaled for desired total time. For analysis, pen position data was low-pass filtered with a second-order Butterworth filter with a cutoff at 10Hz; target positions were converted from screen coordinates (px) to tablet coordinates (mm). The speed-curvature power law was estimated using orthogonal linear regression because both speed and curvature contained measurement uncertainties.

##### Results

For slow, low-frequency movements, the position error (Fig. 2B and Fig. 2E) and velocity errors (Fig 2E) were low, meaning that the participants’ pen position stayed fairly close to target position, and pen speed is also very similar to target speed for all participants. As an example, Fig 2C shows segments of low-frequency tracking speed and curvature time profiles. For each of the target speed profiles (β=0, β=−1/3 and β=−2/3) the participant stays close to the target speed and curvature. However, the speed and curvature of the pen vary more than the speed and curvature of the target. This variance can be seen as the ‘spread’ on the power-law plot (low r^2^). Target log-speed and log-curvature have a strong linear relationship (by design), but participant pen log-speed and log-curvature are uncorrelated.

**Figure 2.**
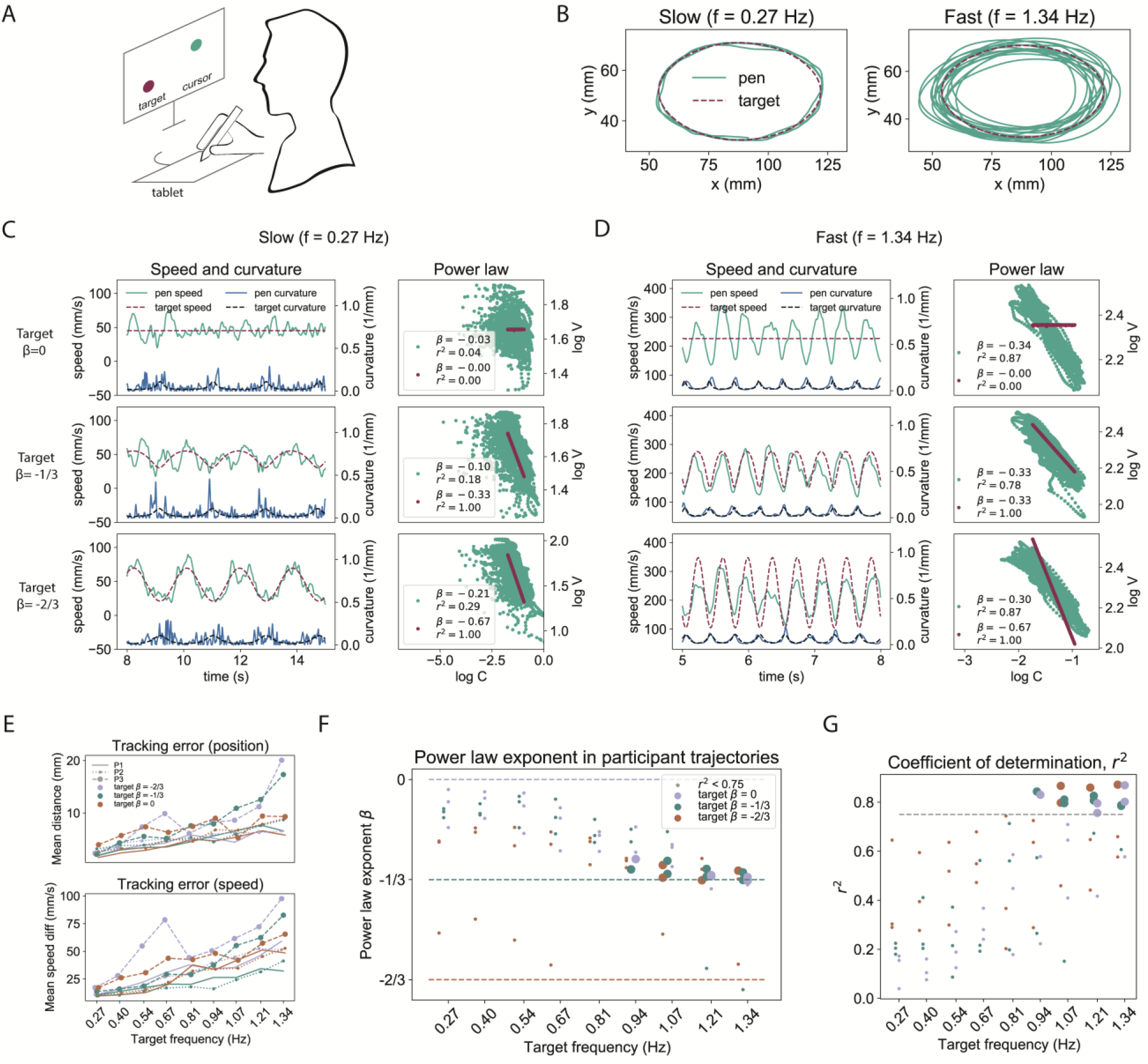
Experiment 1, when tracking elliptic targets, the speed-curvature power law only emerges at high frequencies. **A**) Diagram of the task **B**) Target and pen paths in slow movement (f=0.27Hz, period=3.7s) and in fast movement (f=1.34Hz, period=0.75s). Target ellipse dimensions were 68×39mm, measured from the shape the pen needed to traverse on the tablet. **C**) Segment of the pen and target speed and curvature over time for low-frequency movements, across three different speed profiles; participant’s pen speed seems to track target speed, but pen logC and logV are not strongly correlated **D**) Segment of the pen and target speed and curvature over time for high-frequency movements. The exponent of the pen’s trajectory is β=−1/3, regardless of the exponent of the target’s trajectory. **E**) Participants are more accurate in tracking low frequency targets’ positions and speeds and less accurate in tracking positions and speeds of high-frequency targets, regardless of target speed profile, suggesting poor trajectory control at high frequencies **F**) The speed-curvature power law is only strong enough (r^2^ ≥ 0.75, arbitrary cutoff) for some high frequency movements (f ≥ 0.94Hz), and has the same exponent of β≈−1/3, regardless of target power-law exponent. Each point represents a tracking trial, with the color signifying the target speed profile and size the crossing of threshold of r^2^ **G**) The coefficient of determination (r^2^) rises with the frequency of drawing the ellipses, and crosses the arbitrary threshold of 0.75 at f=0.94Hz.

High-frequency target tracking was very different. There are large errors in position (Fig. 2B, 2E) and large errors in speed (Fig. 2E); however, most participant pen trajectories conform to a speed-curvature power law with the same exponent, β≈−1/3, and a high coefficient of determination r^2^ ≥ 0.75, even though target exponents are different.

An example segment plotted on Figure 2D shows that participant speed had very similar range and mean across different target speed profiles. The power law plots show very similar exponents and high r^2^s in participants’ trajectories, regardless of target exponent. Position and speed tracking accuracy gets worse with the increase in target frequency (Figure 2E), and since the ellipse sizes were always equal, higher frequency meant higher average speed. This result is consistent with the speed-accuracy tradeoff, and suggests poor trajectory control at high average speeds, if trajectory is the controlled variable.

The main results of the first experiment are summarized in Figure 2. Panels F and G: the exponent of the power law of the participants trajectories converges toward β=−1/3 at high frequencies of movement, regardless of target speed profile. The power law was only strong enough (r^2^ ≥ 0.75, arbitrary cutoff) for some trajectories with drawing frequency f ≥ 0.94 Hz (period smaller than ~1.06s), and did not appear for slower movements. The frequency where the power law crosses the threshold of 0.75 also depends on the data filtering procedure, here a second order Butterworth filter with a 10Hz cutoff was used to smooth the position data, and the smoothed data was used estimate curvature and velocity.

#### Experiment 2: Is the cursor-target distance a salient visual feature and a controlled variable?

The instruction to participants in the first experiment was to *keep the cursor as close as possible to the target*. The first guess for the salient visual feature (or visual cue, or visual controlled variable) was the distance between the cursor (C) and the target (T). However, the Euclidean distance is unsigned (always positive), and cannot be used in a proportional feedback control system. In random pursuit tracking it is established that the x and y components of the Euclidean distance (dx and dy) are good approximations for controlled variables (Viviani and Mounoud 1990, Parker et al 2017). In the first test, I’ve analyzed weather dx and dy could be controlled, where dx=Cx – Tx, and dy = Cy – Ty (see the diagram in Figure 2E) in each point in time.

##### Method

For experiment 2, the target trajectory was generated as a smoothed pseudorandom trajectory in two dimensions, and used in a pursuit tracking task (Figure 3A). One participant (male, right-handed) performed the task of following the target with the cursor by moving a pen on an electronic tablet (Huion 610ProV2, recording at 60Hz), with the instructions to keep the cursor as close as possible to the target. The behavior of participants in the ellipse tracking task was first compared to the behavior in the random pursuit tracking task, with the following hypothesis: if the participant controlled the same variables in both tasks, then those variables will have the same relationships to the target position and cursor position, namely dx and dy will be stable and uncorrelated to Tx and Ty, respectively, in both tasks. Next, a numerical model was designed to control dx and dy and then fitted to participant behavior in the pursuit tracking task. The model then performed the ellipse tracking task. If the participants also control dx and dy, their behavior in ellipse tracking should be as well approximated by the model in ellipse tracking task as it is in the random pursuit task.

**Figure 3.**
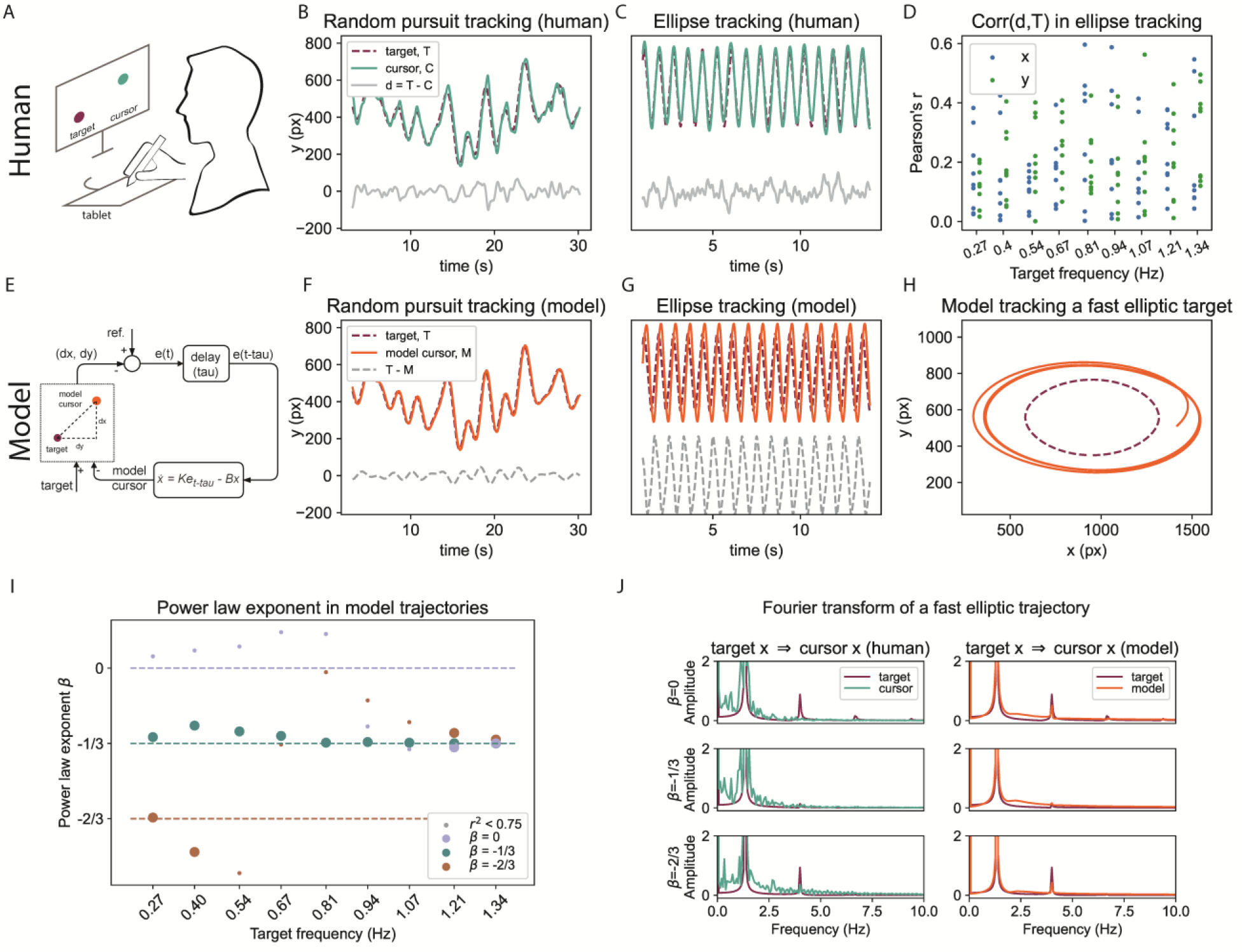
Experiment 2: The cursor-target distances (dx and dy) are controlled variables when tracking pseudorandom targets, but not when tracking elliptic targets. Still, the distance tracking model partially reproduces the β-frequency relationship because of low-pass filtering. **A**) Diagram of the experimental setup. **B**) In a pursuit tracking task, the participant closely follows the target; variables d and T and uncorrelated. **C**) When tracking an elliptic trajectory, the participant is still closely following the target, however, **D**) across the trials in Experiment 1, the correlations between variables dx and Tx, and dy and ty, are not low. **E**) Diagram of a first-order model with delay, parameters fitted in a random pursuit task. **F**) The model controlling the C-T distance follows the target and reproduces participant trajectory in a random pursuit task. **G**) The same model does not account for participant behavior in ellipse tracking, as the dy is larger than in participant’s trajectory and **H**) The elliptic path of the model is larger than the path of the target for fast movements **I**) However, the trajectories of the model converge toward β≈−1/3 power law at high frequencies. **J**) Both the human participant and the numerical model can be seen as low-pass filters of the target signal, passing the low-frequency fundamentals and attenuating the higher-frequency components.

**Figure 4.**
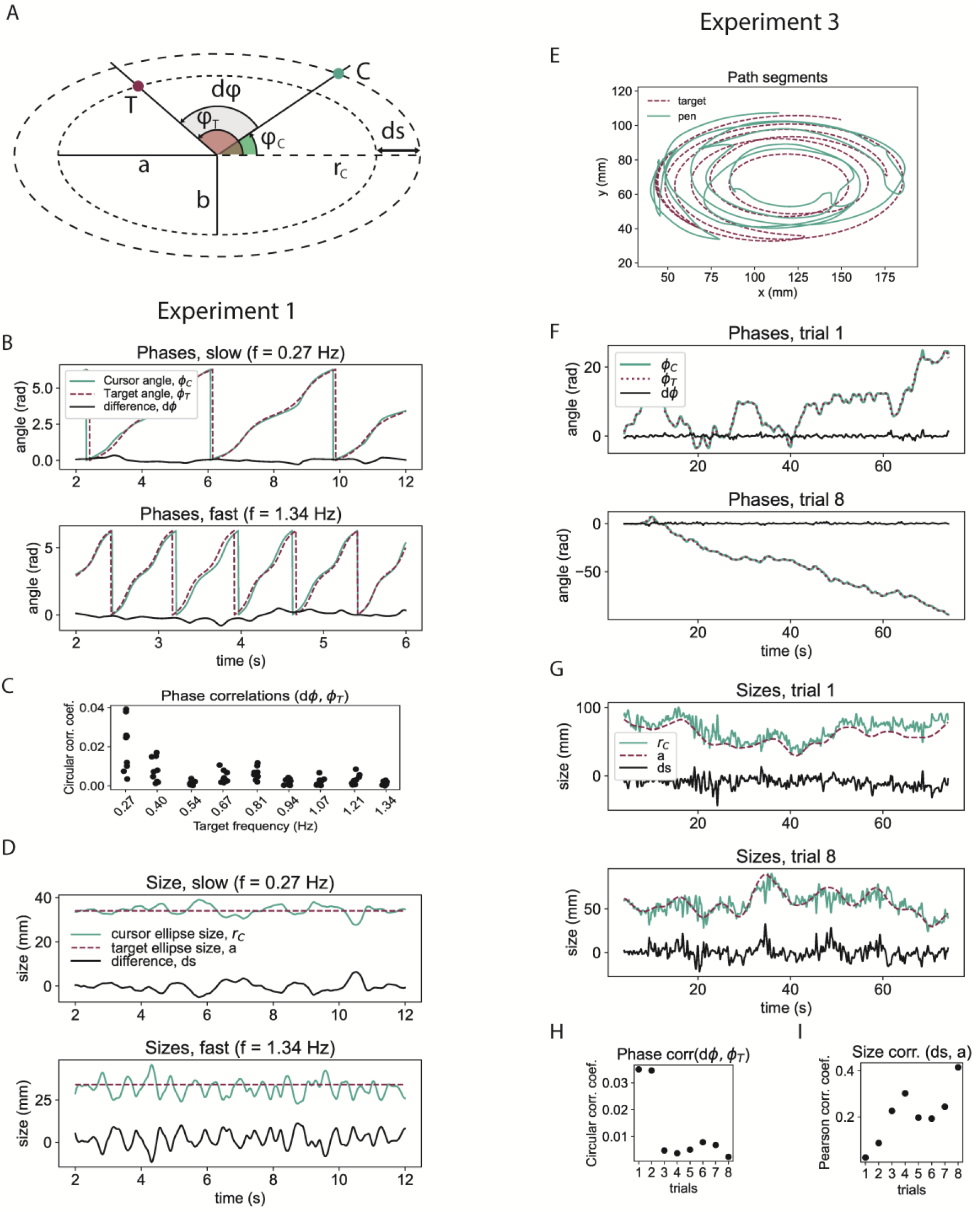
Phase difference (dφ) and size difference (ds) are good candidates for controlled variables in following elliptic trajectory targets. **A**) dφ is the difference between the cursor angle (φ_C_) and target angle (φ_T_) measured from the x-axis; ds is the difference between the x-axis radius of the target path (a), and the x-axis radius of the cursor path (r_C_). The ratio a/b is constant. **B**) Examples from Experiment 1: dφ is stable. **C**) Phase correlations across all tasks in Experiment 1 are very low, using the circular correlation coefficient. **D**) The size of target path (a) is constant, and the size of the cursor path (r_C_) is nearly equal, with low variance. **E**) Experiment 3: cursor and target path segments – target trajectory varies randomly in phase and size. **F**) The participant maintains dφ stable, near zero **G)** The participant maintains ds stable, near zero. **H**) Variables dφ and target angle φ_T_ are uncorrelated **I**) the variables ds and a are only weaky correlated.

##### Results

In the pursuit tracking task, the distance (dx and dy) between the target and the cursor in stable (Figure 3B) and uncorrelated to the target (T), r(dx, Tx)=0.05, r(dy, Ty) = 0.04. Similarly, in tracking elliptic trajectory targets, the distance between the cursor and the target is relatively stable (Figure 3C). However, across all trials, the variables dx and dy are not consistently uncorrelated to Tx and Ty (Figure 3D). This is suggesting there might be better approximations for salient visual features in this task than dx and dy.

In the second line of evidence, a first-order, distance control model with delay (Figure 3E) was fitted to the behavior of the participant. The best fit values for the parameters were found to be: gain K=−10, damping B=0.02 and delay tau=0.100s, similar in x and y. The model accounts for the pursuit tracking behavior fairly well (Figure 3F). However, when the fitted model is tracking the same elliptic trajectory target as the participant, the behavior of the model differs from the behavior of the participant – the distance dy is much larger (Figure 3G), there is phase delay in the model cursor with respect to the target, not seen in participant trajectories, and the amplitude of the model cursor movement is larger than participant amplitude. When the two large-amplitude movements in x and y are combined (Figure 3H), the model cursor trajectory creates a larger elliptic shape than that of the target; while participants maintain the drawn shape on average equal in size to the target-drawn shape.

From these two lines of evidence, we can conclude that dx and dy are not the salient visual features in tracking elliptic targets; they are not controlled in this task, and there might be better approximations for the controlled variables.

While the dx-dy distance model does not account for the behavior of participants in the elliptic target tracking task, model trajectories do show some interesting properties. The exponents of model trajectories, like those of participants, converge to the value of β=−1/3 at high frequencies (Figure 3I). This convergence may be a consequence of low-pass filtering properties of both the model and human participant, as shown by the Fourier transform plot (Figure 3J). Target trajectories that contain harmonics of a frequency higher than the filter’s cutoff frequency will be ‘transformed’ to cursor trajectories that contain only the fundamental frequencies. Similarly, target trajectories with the exponent β=−1/3 across all trials are already single-frequency sinusoids, and when passed through the filter they were phase shifted and amplitude-modulated, they remained single-frequency sinusoids, and their 2D elliptic trajectory conformed to the power law.

#### Phase difference and size difference as controlled variables in tracking elliptic targets

After the confirmation that the cursor-target distances dx and dy are not controlled in tracking targets along elliptic paths, I’ve tested several other hypotheses of controlled variables, and here I report on the most likely ones – phase difference and size difference. The analysis was performed using data from experiment 1.

##### Method

The phase difference (dφ) is here defined as the angle closed by the cursor, the center of the ellipse and the target (Figure 4A), or equivalently, as the difference between the cursor angle (or phase) φ_C_ and the target angle φ_T_. In terms of control systems, dφ is the proposed controlled variable, with the assumed reference value of zero radians, meaning ‘cursor on target’. This controlled variable is disturbed by the changes in the target angle, so the disturbance is φ_T_, and the participant’s behavior acts to bring back the phase difference to zero radians by changing the cursor angle φ_C_. The expectation was that participants maintained the controlled variable stable and uncorrelated to the disturbance variable. Since the variables were measured in radians, the circular coefficient of correlation was used.

The second proposal for the controlled variable is the size difference (ds), defined as the difference between the semimajor axis *a* of the target ellipse (the ‘x-radius’) and the semimajor axis r_C_ of an ellipse with the same center, but passing through the cursor point (Figure 4A). The axis of the cursor ellipse is calculated as 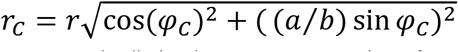, where r is the instantaneous distance of the cursor to the center, and a/b is the constant ratio of target ellipses’ major and minor semiaxes. The value is expected to be stable if it is controlled. However, the correlation coefficient is not defined when one of the variables is a constant (the size of the target ellipse *a*), so the correlation was not calculated.

##### Results

As shown on the figure 4B, the phase difference is maintained stable in the slow trial as well as in a fast trial. Analyzing all the correlations between dφ and φ_T_, for all trials and participants, we can see extremely low values (Figure 4C), supporting the proposal that the phase difference was a controlled variable in the elliptic target tracking task in experiment 1.

Similarly, participants maintained the size difference stable in both slow and fast trials (Figure 4D), supporting the proposal that the size difference was also a controlled variable in experiment 1.

#### Experiment 3: Direct pseudorandom disturbances to phase and size differences

I’ve verified the proposed controlled variables further in a new experiment where the both the size difference and the phase difference were simultaneously disturbed with pseudorandom disturbances, while the participant attempted to maintain them stable. Here, the disturbance to the controlled variable are the target’s angular position and target ellipse size: the participant tracked a target moving along an ellipse in a random-smoothed fashion. Alternatively, the experiment could be performed using the same task as experiment 1 – tracking a target that has a constant rhythm and draws and ellipse of constant size – and the disturbances could be added to the pen position.

##### Method

One participant performed the task of following the target with the cursor by moving the pen on an electronic tablet (Huion 610ProV2, recording at 60Hz), with the instructions to keep the cursor as close as possible to the target. Both the angle of the target and the size of target ellipse varied randomly (Figure 4E). The participant performed 8 trials; the trial number does not signify a progression in any of the task properties.

##### Results

Figure 4E shows a segment of both cursor and target paths in experiment 3. Next, Figure 4F shows participant performance in trials 1 and 8 - the participant maintained the phase difference stable and near zero. The participant also maintained the size difference relatively stable and near zero, or in other words the size of the ellipse shape drawn by the cursor was close to the size of the ellipse shape drawn by the target (Figure 4G).

The correlations of disturbance and the controlled variable are very low in the case of phase angles (Figure 4H), using the circular correlation coefficient. This again indicates that the phase difference is a very good candidate for a controlled variable in tracking elliptic targets. In the case of the size difference, the correlations were moderately low (Figure 4I), using the Pearson’s correlation coefficient, indicating that size difference might be a controlled variable simultaneously with phase difference.

#### Numerical model, sensorimotor phase-locked loop with amplitude control

Having found two good candidates for controlled variables, I’ve constructed a numerical model that controls them, and fitted the parameters of the model to participant behavior in experiment 1. The aim of this model was to reproduce participant behavior in measures such as position, speed, and curvature over time and the exponent β and coefficient of determination r^2^ of the speed-curvature power law. For further validation, the model performed the experiment 3 in the same conditions as the participants, with the same pseudo-random elliptic target trajectories.

##### Method

The model simultaneously controls phase difference and size difference in tracking an elliptically moving target (Figure 5A). It directly incorporates the dx-dy tracking model in the lower level - the loop on the lower right of Figure 5A is the same model seen in Figure 3E. The output of two high-level controllers, phase difference and size difference, is combined to generate the reference signal for the lower-level system. The reference is a point in 2D space acting as a virtual target for the dx-dy tracking system. This control system is aiming to keep the model cursor near the virtual target, using the same controller equation as in experiment 2, and the same parameters obtained when fitting the model to participant behavior.

**Figure 5.**
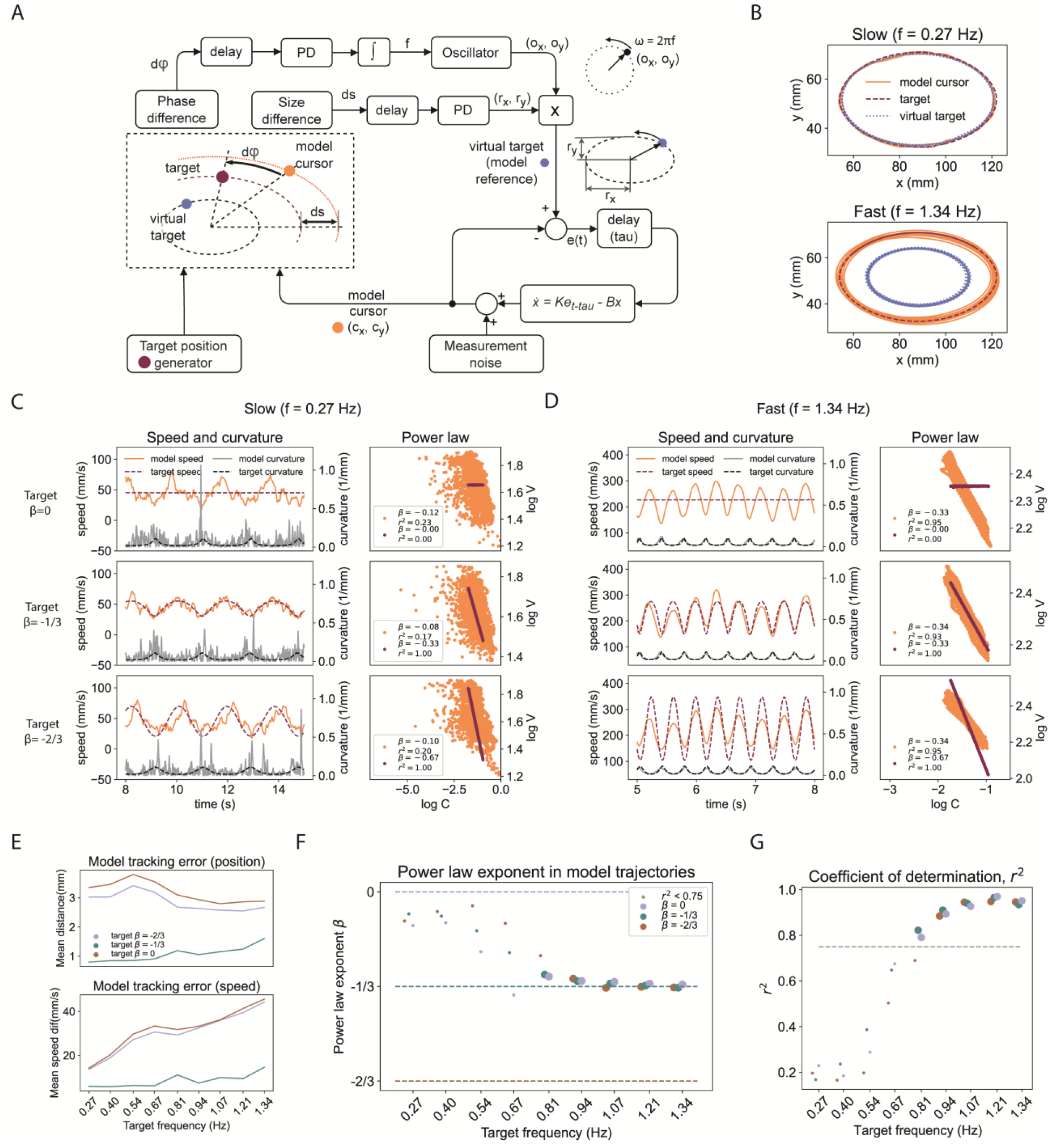
The numerical model reproduces participant trajectories and the dependence of the power law exponent on target frequency found in experiment 1 (Figure 2) **A**) Diagram of the model: a visuomotor phase-locked loop with amplitude control **B**) Model maintains the size of the cursor path equal to the size of the target path for both slow and fast targets by modifying the amplitude of the virtual target **C**) For slow movements, the model’s speed and curvature are similar to participants, including the non-conformity to the speed-curvature power law. **D**) In fast movement, all model cursor trajectories follow the β=−1/3 speed curvature power law, regardless of the target speed profile. **E**) In the given frequency range, the model is much more accurate than participants in both position and speed **F**) The model is reproducing the relationship between the power law exponent β and the movement frequency found in Experiment 1, as well as **G**) the relationship between the coefficient of determination r^2^ and movement frequency.

The phase control loop in this model is a version of the phase-locked-loop (PLL), a very common control system in radio communication and information technology used to synchronize phases and frequencies of signals. Keeping the phases of two signals in ‘lock’, where their difference is a constant value (not necessarily zero), means that their frequencies will be equal.

The main parts of the phase control loop appear on the diagram (Figure 5A) starting with the phase difference ‘detector’. Given positions of the center of the ellipse, the target and the cursor, the phase difference detector finds dφ: the angles of φ_T_ and φ_C_ are calculated as arctangents of the x and y components of their positions relative to the center and unwrapped. The output of the phase difference detector is dφ = φ_T_ − φ_C_. The value dφ is passed to a pure delay element and used as an input to a proportional-derivative controller (PD) with an implicit reference of zero, so the phase difference is maintained at zero. The error is integrated as passed to the harmonic oscillator as the frequency parameter f. The oscillator is implemented as a rotating unit vector with instantaneous frequency f and angular velocity ω=2πf, described by the equations 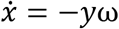 and 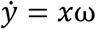 with initial angle set to the target angle φ_T_.

The size difference control loop maintains the size of the cursor-drawn ellipse equal to the size of the target drawn ellipse. The size r_C_ is calculated as 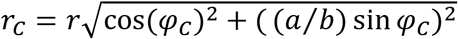, where r is the instantaneous distance of the modelled cursor to the center, and a/b is the constant ratio of target ellipses’ major and minor semiaxes. The controlled variable ds is the difference between the target ellipse size determined by the parameter **a**, and the size of the cursor ellipse r_C_. The difference is passed to a pure delay element, then a PD controller gives values rx and ry, where ry = (a/b) rx. The values rx and ry are multiplied with oscillator output to generate the 2D location of the virtual target.

The lower-level loop aims to maintain the cursor on the virtual target. It is identical to the dx and dy control loop from experiment 2, has the same parameters and therefore identical low-pass filtering properties, presumably matching those of the participant. The difference is that the ‘target’ is an internal reference signal, and not an external stimulus. In experiment 2, we observed that the dx and dy control model lagged in phase behind the target, and that the size of the ellipse drawn by the model was different than the size of the ellipse drawn by the target. The role of the virtual target was to automatically compensate for the attenuation or amplification of amplitude produced by the low-pass filtering process. The size of the ellipse drawn by the virtual target was automatically increased when there was a size difference between the cursor and the ‘real’ target. Similarly, the virtual target was automatically advanced in phase to compensate for the phase lag produced by the low pass filter.

Another important part of this model is the simulated measurement noise. This a is a small-amplitude, normally distributed pseudorandom signal (mean=0mm, std=0.06 mm), added to cursor in x and y position independently during the task. This signal was primarily aimed at modelling the noise arising in the electronic tablet and pen instruments during recording.

The model performed the same task as the participants of first tracking the target across elliptic trajectories from experiment 1, and second across randomly varying phase and size from experiment 2. Recorded position trajectories were smoothed with a 10Hz cutoff low-pass, second-order Butterworth filter. The simulation step was 5ms, using Euler integration. Time derivatives of signals were approximated as differences between current and previous time-step signal values, divided by the time-step length.

##### Results

The parameters of the model – the gains and delays of the phase and size difference control loops – were found by fitting the behavior the model to the behavior of one participant. The parameters of the inner loop were found in experiment 2, and they remained the same (K=10, B=0.018 and delay=100ms). The size difference loop had a delay of 250ms: 150ms on top of 100ms from the inner loop, while the phase difference had a delay of 100ms; no additional delay on top of the inner loop. The phase difference proportional-derivative (PD) controller had Kp=0.5 and Kd=0.3, while the size difference PD controller had Kp=1.33 and Kd=0.016, and an additional slowing term with B=0.033.

Figure 5 shows the performance of the model in the same tracking task performed by the participants, analyzed in the same way as participant performance (Figure 2), while Figure 6 shows a direct comparison between one participant and the model.

**Figure 6.**
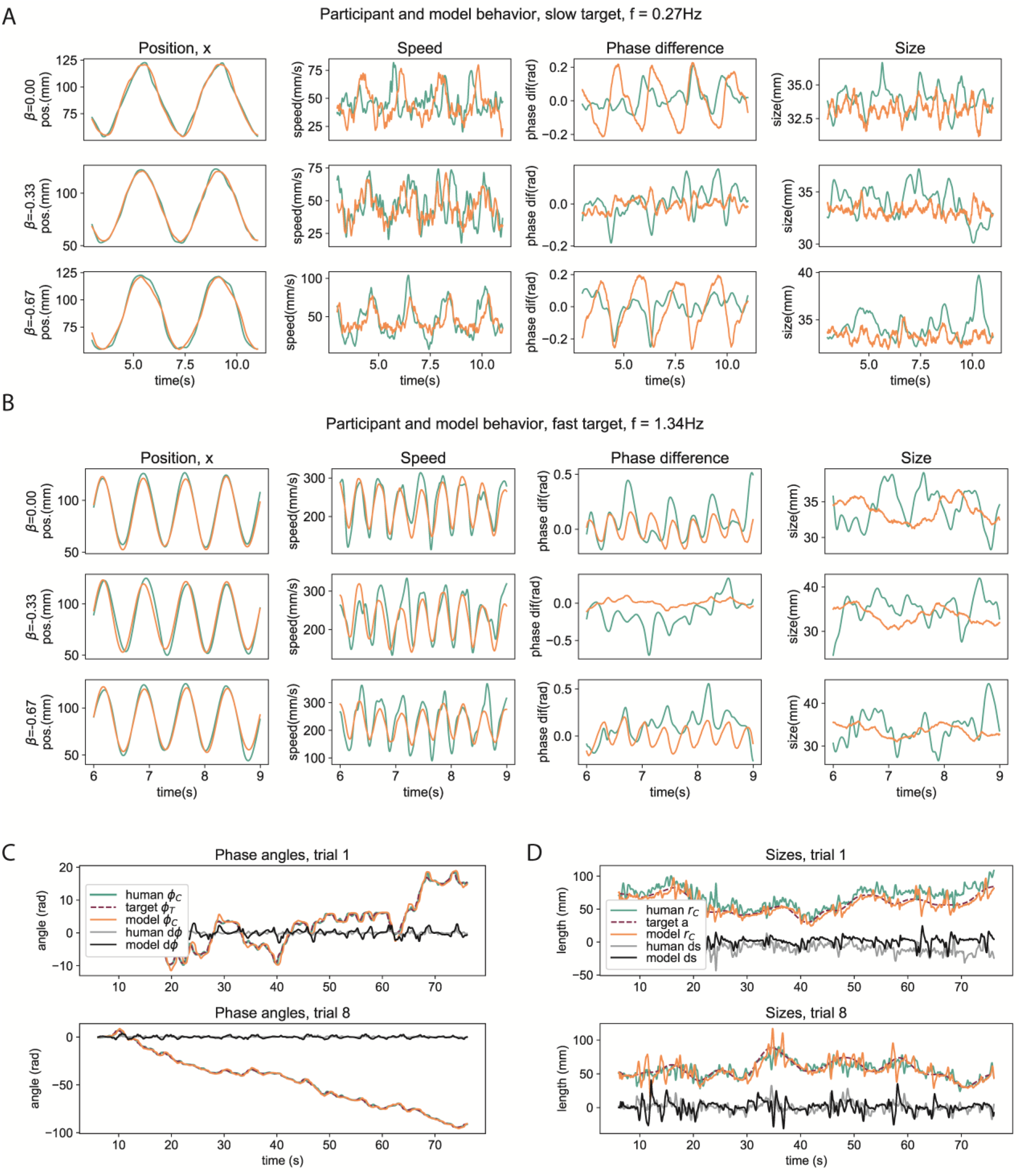
Direct comparison of one participant and model behavior in the same tasks (model in orange, participant in green). **A**) In tracking a low frequency (slow) target, model and participant behave similarly in measures of position in the x dimension, speed, phase difference and ellipse size. **B**) In tracking a high-frequency (fast) target, the model shows more regularity, while the participant shows more randomness in behavior, however the positions and speeds are very similar **C**) The model also reproduces participant phase angles from Experiment 3, where the target phase varied randomly **D**) The model reproduces participant ellipse sizes from Experiment 3, where the target size varied randomly.

The model closely approximates the behavior of the participants – the speed curvature power law emerges only for high-frequency target tracking, where r^2^ ≥ 0.75 only for trials where the target frequency was f ≥ 0.81Hz (Figure 5F and 5G), while for participants the power law was present for f ≥ 0.94Hz.

In a slow trial (target f=0.27 Hz, period=3.7s, Figure 5C), for different target speed profiles (β=0, β=−1/3, β=−2/3), the model cursor seems to be tracking the low-frequency fundamental, and there is some higher-frequency noise, in both speed and curvature profiles. Speed and curvature are only weakly related and the power law does not reach r^2^ ≥ 0.75. This is similar to participant speed and curvature profiles for the same task - compare Fig. 5C to Fig. 2C.

In a fast trial (target f=1.34Hz, period=0.75s, Figure 5D), for different target speed profiles, the model has the same speed profile, with the exponent β≈−1/3, and with high r^2^. This is similar to the participant speed profiles, and the relationship between speed and curvature in participant trajectories – compare Fig 5D to Fig 2D.

The position and speed errors across trials with different frequencies were not the same as participants (Figure 5E, compare to Figure 2E). The model has lower errors, lowest when the target speed profile conforms to β=−1/3 power law. This distinction does not appear strongly in participant position and speed errors plot (Figure 2E). The absolute value of the errors is also smaller in the model than in participant trajectories. In the given range of target frequencies, the position error for targets with β=0 and β=−2/3 is relatively large even for slow targets, and does not have a clear trend.

The similarity of the behavior of the model is to the behavior of the participants is best seen in direct comparison of several time profiles: position, speed, phase difference and size (Figure 6). In the slow, low frequency trial (Figure 6A) the position of the model closely matched participant position in the same task. Speed profiles contain similar amounts of high-frequency components, mostly coming from the modelled low-amplitude measurement noise.

In phase difference and size, the participant’s profiles seem to contain more randomness, while model phase difference and size are more repetitive. Direct comparison between the participant and the model in fast movement (Figure 6B) reveals very similar position and speed profiles, while phase difference and size seem to be more variable in participants trajectories than in the model trajectories.

Without changing the parameters, the model performed a tracking trial identical to experiment 3 (Figure 6C and 6D). The model maintained simultaneously the phase difference (Figure 6C) and the size difference (Figure 6C) stable and near zero, in a very similar range to that of participants, even when both target phase and ellipse sizes had a randomly-varying time profile.

## Discussion

### The power law does not necessarily appear for low-speed trajectories

The main empirical finding in experiment 1 (Figure 2) was that the speed-curvature power law appeared (r^2^ ≥0.75) only for high frequency movements, when the fundamental frequency was larger than 0.94Hz, or equivalently, when the period of drawing an ellipse was shorter than 1.04 seconds, and average speed higher than 158mm/s. For higher rhythms, participants did not track instantaneous target speeds well. Instead, participants’ speed profiles were nearly the same from trial to trial, following the β=−1/3 speed-curvature power law, regardless of the target’s power law exponent. In other words, for fast rhythms, participants always moved slightly slower in the curves than in the straight parts, even when the target moved at a constant speed (β=0) or had a higher speed difference profile (β=−2/3) (Figure 2D). In contrast, for low frequencies, (low average speed, cycle periods longer than 1.04s) the fit to the speed-curvature power law was poor. Participants roughly tracked instantaneous positions and speeds of the targets (Figure 2E) but the curvature and speed profiles contained a lot of high-frequency components, possibly coming from movement noise and measurement noise.

The limit where the power law fit (r^2^) reaches 0.75, at period time of 1 second, or frequency of 1Hz, might be specific to small-size elliptic trajectories of the hand, since the target ellipse had a constant size, 6.8cm in width and 3.7 cm in height. It has already been reported that drawing larger elliptic shapes does not always result in a good fit to the power law (Schaal and Sternad, 2001). Future research should look into the interaction of average speed, size of the trajectory and rhythm or period of drawing, and their influence on the exponent and the strength of the speed-curvature power law.

At low speeds, the participants did closely reproduce the low-frequency target trajectories in both speed and position (Figure 2E). If participants can explicitly track or control trajectories, this ability seems to have a fairly low bandwidth, as the errors steadily increase with frequency, and yet, the size of the elliptic shape and the rhythmic synchronization to the target were still preserved. This seems to suggest a non-trivial control scheme, where *trajectory is not the controlled variable*.

### Cursor-target lineal distance is not controlled

In experiment 1, participants were instructed to *‘keep the cursor as close to the target as possible’*, and the first approximation for the controlled variables were the x and y components of the cursor-target distance, variables called dx and dy. There are several independent lines of evidence suggesting dx and dy are *not* controlled in ellipse tracking. First, in random pursuit tracking, dx and dy are demonstrably good approximations for controlled variables (Parker, 2017), and as expected, there are very low correlations between dx and Tx, as well as between dy and Ty in experiment 2 (random pursuit, Figure 3B). However, those correlations were higher in ellipse tracking, suggesting that dx and dy are not controlled in ellipse tracking (Figure 3D).

Second, the dx-dy model made much bigger ellipse drawings than the participants, and created a large phase lag, again suggesting that dx and dy are not controlled in ellipse tracking.

### Low-pass filtering accounts for the power law

The low-pass filtering properties of the dx-dy model may account for the convergence of the power-law exponents toward a value of β=−1/3 at high speeds (Figure 3I and 3J). A similar result was found by Gribble and Ostry (1996), where they concluded that the power law may arise from the low-pass filtering properties of the musculoskeletal system of the arm. Schaal and Sternad (2001) found that even a simple Butterworth filter with an appropriate cutoff frequency can produce power-law trajectories out of constant-speed trajectories. Recently, we have shown that a robot arm following a constant-speed visual target can also produce power law trajectories because it behaves as second order system with delay, also a low-pass filter (Matić et al., 2021). Similarly, as recognized by Lacquanity et al (1983), an ellipse composed of pure sinewave components, without any harmonics, will conform to the −1/3 speed-curvature power law. Low-pass filtering creates smooth trajectories for frequencies below the cutoff, and this alone might explain the speed-curvature relationship – if the target trajectory is the input signal, the visuomotor and proprioceptive-motor loops are filters that smooth it out, and create pure sinewaves in the cursor (or pen) trajectory as output.

The difficulty with this explanation is that low-pass filters distort the input signal by introducing amplitude modulation and phase lags – filtered trajectories in Schaal and Sternad (2001), Gribble and Ostry (2003) and Matić et al (2021) all show that output ellipses have a different size than input ellipses. Participants in experiment 1 generally maintained the size of the cursor trajectory equal to that of the target’s trajectory, and followed the targets without phase lags – similar results were found by Viviani and Mounoud (1990) when tracking ellipses and by Parker (2020) when tracking one-dimensional sine waves.

A possible solution is that higher-order systems somehow compensate for the distortions introduced by the filter.

### Phase and size difference may be controlled

I’ve proposed that higher order systems directly observe and control the visual phase difference and the size difference between the ellipses drawn by the cursor and the target. In experiment 1, the phase difference was maintained stable, and also had a very low correlation with disturbances (Figure 4B, 4C), which means it was unaffected by the disturbances and likely controlled. Participants also maintained the size of the cursor-drawn ellipse relatively constant (Figure 4D), which means that the size difference was also likely controlled.

In experiment 3, direct disturbances were applied to the phase difference and size difference variables by designing an elliptic target trajectory that had a randomly changing phase and size. The stability and correlation measures confirmed the findings from the first experiment (Figure 4E, 4F, 4G, 4I).

Instead of creating new target trajectories, an alternative strategy might be to apply random perturbations in the pathway between the pen and the cursor, making the disturbances invisible to the participants, and maintaining task visually equal to experiment 1, but altering the pen movement patterns necessary to maintain phase difference and size difference stable.

### Numerical model reproduces participant trajectories

For the next verification step, I’ve created a numerical model, a simulation of the perceptual and control processes of participants. It is implemented as a two-level hierarchical control system that perceives and controls the cursor-target phase difference and the cursor-target ellipse size difference (Figure 5A). The phase difference is integrated in time and used as a frequency parameter to a harmonic oscillator. The output of the oscillator is multiplied by the size difference error to create a virtual target – a phase-advanced and amplitude-corrected reference for the inner loop. When the model cursor follows the virtual target, the phase difference and size difference can be kept low in the whole range of target frequencies tested; making the behavior of the model similar to human participants. The low-pass filtering properties of the dx-dy model are maintained because the inner loop has the same structure and parameters.

The model replicated some important measures of participant behavior – namely the absence of the speed-curvature power law at low frequencies and the emergence of the power law at high frequencies, with the same exponent β≈−1/3, and r^2^ ≥ 0.75 for higher frequencies (f ≥ 0.84Hz). For participants this frequency was f ≥ 0.94Hz, and the difference is possibly a consequence of unmodelled, multiplicative noise (Faisal et al., 2008). Speed and position profiles of the model cursor were also similar to participants (Figure 6). On the other hand, the speed and position errors (the distance between the cursor and the target) in the model (Figure 5E) were lower than the speed and position errors in participant trajectories (Figure 2E) - the model was more accurate. This might suggest some changes in the model that would make it less accurate in speed and position tracking, while still maintaining a good fit in other measures.

The model also replicated participant behavior in non-rhythmical target trajectories, namely when both the phase of the target and size of the target ellipse had pseudorandom profiles (experiment 3, Figure 6D and 6E), suggesting that the model might be more general – the target trajectory does not need to be rhythmical, and the size of the target ellipse does not need to be constant. The model performed all the tasks without any changes in the parameter after initial fitting.

### Parameter measurement implications

If model parameters represent some internal characteristics of the participant, their values might be informative about the characteristics of the participant. The delay parameter represents the time it takes for a signal to make a ‘full trip’ around the loop, and longer delays might indicate longer computational processing or longer nerve pathways, implying hierarchically higher systems. As expected for a hierarchically higher system - the size difference control loop has a large delay (250ms), but the phase difference loop is only at 100ms, the same as the inner control loop. This might indicate that the model needs some reorganizing. I’ve proposed the phase control loop as superordinate to the inner cursor position loop, but those two loops might also be on the same organizational level. The proposed oscillator is ‘cortical’, since it sends outputs downstream to the visual system, however it might be a spinal or a brainstem central pattern generator. This proposal could be verified in a future study.

### Alternative and additional controlled variables

As mentioned in the introduction, there are always potential improvements to the approximation of the controlled variables. Similar measures to phase and size difference might also be used, such as the arc-length distance instead of the phase difference. This might be a more useful measure when the target path is not elliptical. The size of the ellipse, as proposed, might be split into the horizontal and vertical amplitude components, allowing the control of elliptic paths of independently varying width and height. Alternatively, path accuracy might be controlled more locally, by perceiving the distance between the cursor and the path, and altering the direction of movement. The velocity of the target itself might play a role, as in one-dimensional sinewave tracking (Parker et al., 2021). An additional control variable in the task might be the center of the oscillation.

### Limitations of the study

Participants were not trained to perform the task. The task was not difficult, however, training until there is no more improvement should remove the influence of learning on performance and make the participant trajectories less variable from cycle to cycle.

There were only three participants in the first experiment, and a single participant in the second and third. A larger number of participants who would perform all three experiments would allow fitting the model to each participant individually, and then testing how well the model predicts their behavior in a new task. While I have not used any statistical inference tests that would require a large number of participants, a higher number of participants might still be useful to gather the ranges and consistency measures of model parameters, such as gains and delays, in the sample.

### Summary and outcome

In this study, I’ve attempted to answer why there is a relationship between speed and curvature in elliptic hand movement, why is the exponent of the speed-curvature power law often found to be −1/3, and more broadly – how people track elliptic trajectories.

I’ve found that the power law only appears in fast movement, here when elliptic paths are traversed in less than 1 second, for relatively small elliptic shapes (6.8 cm width and 3.7 cm height).

Based on the recorded data, I’ve proposed phase difference and size difference as controlled variables in the ellipse tracking task and supported the hypothesis by finding that participants can maintain them stable when they are directly perturbed in a tracking task. Also, there are low correlations between controlled variables and disturbance variables in two different tasks.

Finally, a numerical model that controlled phase and size difference reproduced many of the behavioral measures found in participants’ hand trajectories, such as the emergence of the speed-curvature power law, as well as position and speed profiles of participants’ hand trajectories.

In conclusion, it appears that in ellipse tracking, the speed-curvature power law emerges because of the low-pass filtering properties of the whole visuomotor loop, including visual processing and musculoskeletal elements. The effects of the low-pass filter – phase delay and amplitude modulation – are compensated by two higher level loops that maintain the phase of the cursor equal to the target and the size of the drawn ellipse equal to the size of the target ellipse by modulating the frequency and amplitude of a simple harmonic oscillator.

## Acknowledgements

I would like to thank Alex Gomez-Marin for his involvement in the design of Experiment 1, and the comments of the figures.

## Notes

https://github.com/adam-matic/visuomotor-phase-locked-loop

### Competing Interest Statement

The authors have declared no competing interest.

### Summary of Updates

Higher resolution figures and minor edits or typos. Added acknowledgements.

